# Mapping general anesthesia states based on electro-encephalogram transition phases

**DOI:** 10.1101/2023.07.06.547567

**Authors:** V. Loison, Y. Voskobiynyk, B. Lindquist, D. Necula, D. Longrois, J. Paz, D. Holcman

## Abstract

Cortical electro-encephalography (EEG) has become the clinical reference for monitoring unconsciousness during general anesthesia. The current EEG-based monitors classify general anesthesia states simply as underdosed, adequate, or overdosed, with no transition phases among these states, and therefore no predictive power. To address the issue of transition phases, we analyzed EEG signal of isoflurane-induced general anesthesia in mice. We adopted a data-driven approach and utilized signal processing to track *θ*- and *δ*- band dynamics as well as iso-electric suppressions. By combining this approach with machine learning, we developed a fully-automated algorithm. We found that the dampening of the *δ*-band occurred several minutes before significant iso-electric suppression episodes. Additionally, we observed a distinct *γ*-frequency oscillation that persisted for several minutes during the recovery phase following isoflurane-induced overdose. Finally, we constructed a map summarizing multiple states and their transitions which can be utilized to predict and prevent overdose during general anesthesia. The transition phases we identified and algorithm we developed may allow clinicians to prevent inadequate anesthesia, and thus individually tailor anesthetic regimens.

**Significance statement:** In human patients, overdosing during general anesthesia can lead to cognitive impairment. Cortical electro-encephalograms are used to measure the depth of anesthesia. They allow for correction, but not prevention, of overdose. However, data-driven approaches open new possibilities to predict the depth of anesthesia. We established an electro-encephalogram signalprocessing pipeline, and constructed a predictive map representing an ensemble of gradual sedation states during general anesthesia in mice. In particular, we identified key electroencephalogram patterns which anticipate signs of overdose several minutes before they occur. Our results bring a novel paradigm to the medical community, allowing for the development of individually tailored and predictive anesthetic regimens.

## 2 Introduction

A century of analysis of cortical electro-encephalogram (EEG) data has provided comprehensive classifications of brain states. In the last 50 years, spectral analysis and signal segmentation of the EEG have matured enough to provide relevant information about the instantaneous dynamics of brain activity during sleep [1], coma [2, 3], and general anesthesia (GA) [4, 5]. EEG is routinely used to monitor the adequacy, also known as depth, of anesthesia in humans. Indeed, although recent data demonstrate associations between anesthetic overdose and post-operative complications [6, 7], whether real-time monitoring overdose alerts provided to anesthesiologists improve outcome is still controversial [8, 9, 10]. These observations suggest that prevention of hypnotic overdose is necessary in order to improve outcome. Prevention of hypnotics overdose, as opposed to correction after the fact, would require a new paradigm for EEG analysis.

When patients are anesthetized with propofol or halogenated gas such as sevoflurane or isoflurane, the brain transitions to a state characterized by the presence of frontal *α*-oscillations in the 8 – 12 Hz range [11]. When the hypnotic concentration increases, the *α*-oscillations can disappear, leading to partial suppressions of the *α*-band known as *α*-suppressions (*α*S) [12, 13]. In contrast, in rodents, GA is characterized by the presence of *θ*- and *δ*- oscillations [14]. Further increase in hypnotic concentration in humans and rodents can lead to periods of flat EEG, called iso-electric suppressions (IES). Occurrence of IES marks profound anesthesia that has been robustly associated with post-anesthetic complications in humans, such as delirium and cognitive dysfunction [15, 16]. Post-operative delirium has also been observed in mice [17], but its link to IES has not been rigorously investigated. Interestingly, temporal relationships have been established between EEG patterns in humans. For instance, suppressions of the *α*-band precede IES appearance in humans anesthetized with propofol [12]. It is thus possible to predict which patient will be most sensitive to overdose from the first 10 minutes of GA with propofol [13]. However, no similar approach exists for GA obtained with halogenated gas. Indeed, propofol and halogenated gas induce different EEG signatures [18]. We thus decided to investigate whether temporal relationships between EEG patterns could be found in isoflurane-induced GA.

Several computational methods are commonly used to analyze EEG, including wavelets [19, 20], thresholding methods [21], and empirical mode decomposition [22]. Based on the EEG and electro-myogram (EMG) recordings of isoflurane-induced GA in mice, we developed a novel signal-processing approach, coupled with machine learning. This coupling allowed us to discover relevant patterns and evaluate their predictive power. The present EEG time-frequency analysis relies on the IRASA decomposition algorithm [23] which isolates oscillatory components from background spectral decay. By computing the relative power of frequency bands and the time proportion of IES, we uncovered multiple EEG and EMG states and found specific deterministic transitions between them. This led us to propose a state chart to represent brain states and their associated transitions. This state chart could be used to assess and predict depth of anesthesia in isoflurane-induced GA.

## 3 Results

### 3.1 Higher isoflurane concentration increases IES incidence

To evaluate whether the GA protocol could impact the appearance and distribution of IES, we implemented three different protocols with varying isoflurane concentrations. In the first, which we refer to as the incremented protocol, the isoflurane concentration was gradually increased from 0.5% to 2% every 5 minutes in 0.5% increments (n=13). In the other protocols the isoflurane concentration was fixed at 1% (n=9), and 1.5% (n=8) (Fig. 1A-B). We refer to these protocols as the 1% protocol and the 1.5% protocol respectively.

**Figure 1:**
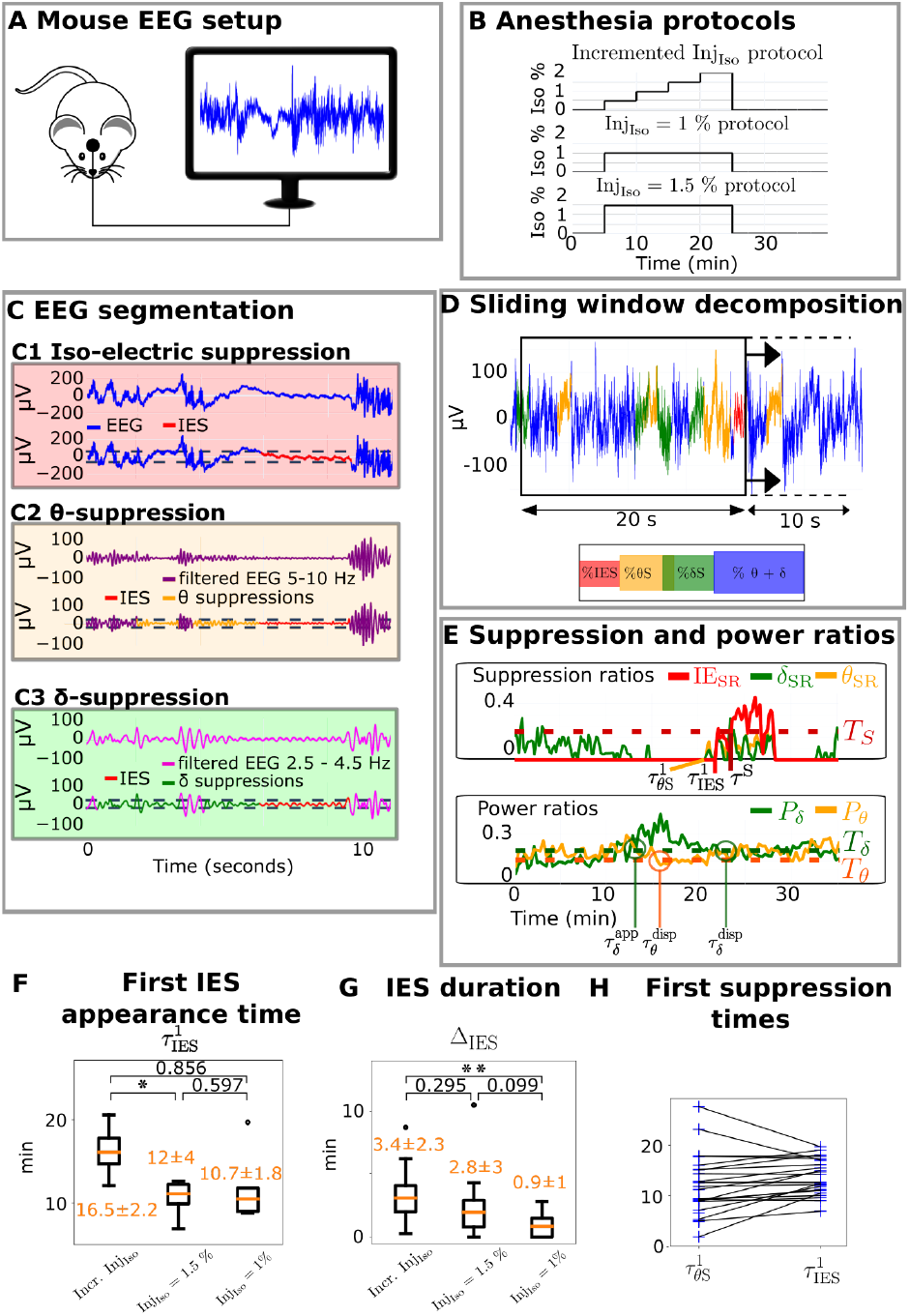
Time-frequency segmentation of EEG recorded during isoflurane-induced GA in mice. **(A)** Experimental setting for a single-electrode EEG recording. **(B)** Three protocols used for isoflurane-induced GA. **(C)** Suppression detection leading to **(C1)** IES (red), **(C2)** *θ*-suppression (yellow), **(C3)** *δ*-suppression (green). Segments where the signal is lower than a threshold (dotted lines) are labeled as suppressions (see Methods). **(D)** Time spent in each suppression type in a sliding time window. **(E)** Suppression ratios across the EEG recording, with the delay to first IES 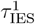, the IES ratio exceeds threshold *T*_*S*_ at time *τ* ^*S*^, and relative power of the *θ* and *δ* rhythms. **(F)** Distribution of delay from GA induction to first IES 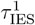. **(G)** Distribution of cumulative time spent in IES Δ_IES_. **(H)** Delay from GA induction to the first *θ*-suppression occurrence, and first IES occurrence respectively. The black lines link two points coming from the same recording. \**P <*0.05, \*\**P <* 0.001, two-sided Wilcoxon-rank U test.

We used an automated algorithm adapted from [12] to detect IES and suppressions of the prominent bands, namely the *θ*-band (4*−*10 Hz) and the *δ*-band (0*−*4 Hz) (Fig. 1C and Methods). We computed the suppression ratios IES_SR_, *θ*_SR_ and *δ*_SR_, which we defined as the duration ratio spent in iso-electric suppression, *θ*-suppression, and *δ*-suppression respectively, over a 20-second sliding time window (Fig. 1D and Methods). We then identified the delay 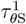 to the first *θ*-suppression occurrence, the delay 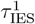 to the first IES occurrence, and the delay to start of long IES *τ* ^*S*^ (defined as the time at which the iso-electric suppression ratio IES_SR_ exceeds the empirically chosen threshold *T*_*S*_=0.25) (Fig. 1E and Methods). Using the same sliding-window analysis, we also computed the time-dependent relative powers of the *δ*- and *θ*- bands (see Methods).

For the incremented protocol, IES appeared uniformly in 100% (13/13) of mice, 16.5 ± 2.2 minutes after GA onset, at 2% isoflurane concentration. When using 1.5% concentration in a subset of the same mice, IES appeared in 89% (8/9) of mice, 10.7 ± 1.8 minutes after GA onset. However, at 1% isoflurane concentration, our algorithm detected IES in only 62.5% (5/8) of mice, 12 *±* 4 minutes after GA onset (Fig. 1F). We found a statistical difference in the IES duration Δ_IES_ (defined as the cumulative time spent in IES per recording) between the incremental protocol and the 1% protocol but not between other protocols (Fig. 1G).

Interestingly, the *θ*-band was suppressed at the same time as IES, but not before (Fig. 1C2, Fig. 1H), contrasting with propofol-induced GA in humans, where frontal *α*-suppressions precede IES [12, 13]. We, therefore, conclude that suppressions of the *θ*- and *δ*-bands do not reliably precede IES appearance in parietal EEG recordings from mice during isoflurane-induced GA.

### 3.2 Spectral decomposition reveals *θ*- and *δ*- bands predominance

To analyze the EEG recordings, we proceeded with a spectral approach. We started with a decomposition on a single time window *W*_*t*_ (Fig. 2A). For each window *W*_*t*_, the power spectral density was decomposed into two components: the first was the 1/f-component that accounted for a decaying trend, fitted by the function 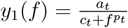 (Fig. 2A2, see Methods). The second was the oscillatory component (Fig. 2A3) accounting for the activity in isolated frequency bands, which we fitted with Gaussians (Fig. 2A4). This separation was extended to the entire recording by sliding the time window *W*_*t*_ (Fig. 2B-D). Our algorithm faithfully separated the oscillatory component and tracked it continuously over time (Fig. 3D, see Methods). We found that in isoflurane-induced GA in mice, two frequency bands were active: the *θ*- and *δ*-rhythm (illustrated in Fig. 3D black lines and pink lines, respectively). The *θ*-rhythm was always present before GA induction and decayed shortly after induction. The *δ*-rhythm appeared few minutes after GA induction.

**Figure 2:**
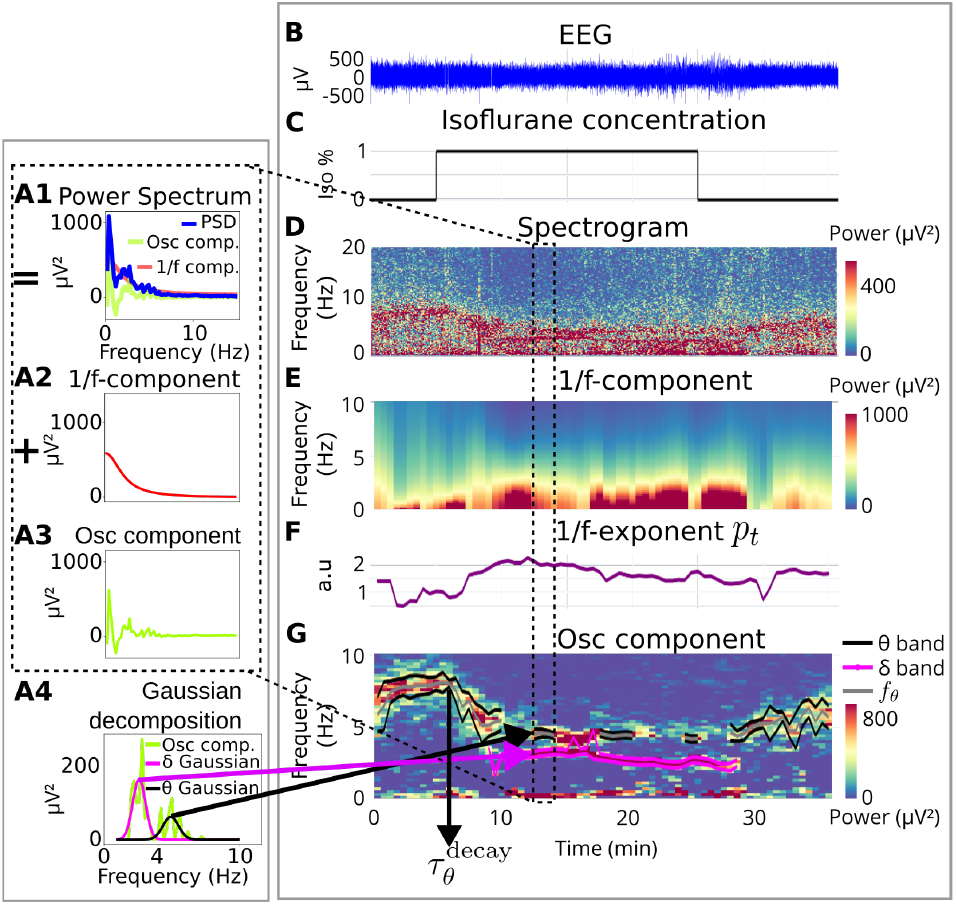
EEG Spectral analysis applied to isoflurane-induced GA in mice on a single one-minute time window **(A)**, and along the entire procedure **(B-G)**. The Power Spectral Density (PSD) of one minute of EEG is computed **(A1)**, and separated into the 1/f-component **(A2)** and the oscillatory component **(A3)** (eq. 12). The *θ*- (pink) and *δ*- (black) band characteristics of this time window are obtained by fitting Gaussians to the oscillatory component **(A4)**. This separation is performed on successive time windows along the procedure, resulting in a continuous estimation of the spectral parameters. **(B)** EEG recording on which the estimation is applied. **(C)** Anesthesia protocol. **(D)** Spectrogram of the entire EEG recording. **(E)** Estimated 1/f component. **(F)** Exponent of 1/f component. **(G)** Estimated oscillatory component and tracked *θ* and *δ* rhythms.

**Figure 3:**
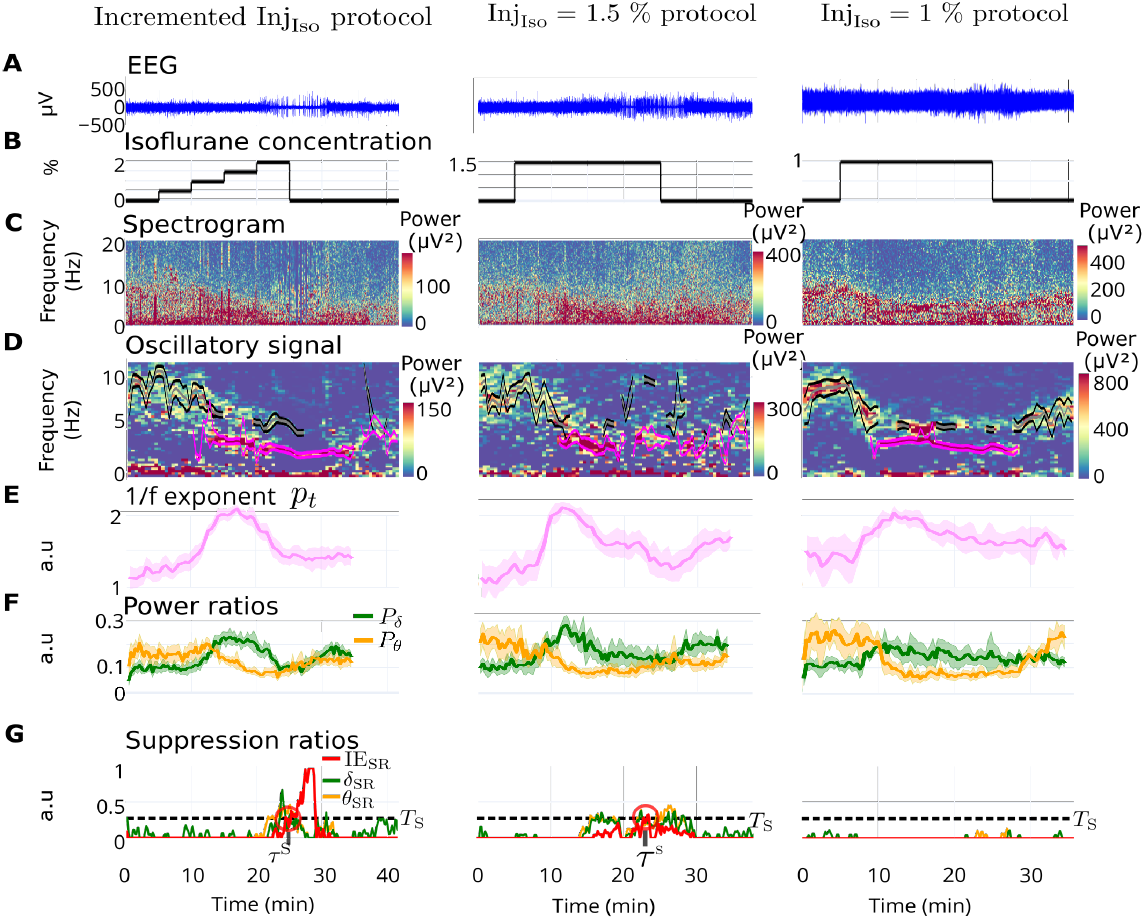
Influence of anesthesia protocols on spectral dynamics. **(A)** Representative EEG examples of three GA protocols **(B). (C)** Spectrograms.**(D)** Oscillatory components and automatically tracked *θ*- (grey and black) and *δ*- (pink) rhythms. **(E)** Average exponent *p*_*t*_ of the 1/f-component 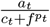 per protocol. **(F)** Average power ratios *P*_*δ*_ (green) and *P*_*θ*_ (yellow) per protocol. **(G)** Examples of suppression ratios computed for the *δ*-band (*δ*_SR_), *θ*-band (*θ*_SR_) and the IES (IES_SR_). Error bands indicate the 95% confidence intervals computed using the *t*-distribution.

Interestingly, the dynamics of the 1/f exponent *p*_*t*_ revealed a general trend across all recordings and protocols: *p*_*t*_ increased during the beginning of GA, then reached a plateau and decreased (Fig. 3E and supplementary section S1.1).

Notably, the relative power of the *θ*-band *P*_*θ*_ (yellow) was reliably higher than the relative *δ* power *P*_*δ*_ (green) before the beginning of GA (Fig. 3F). Shortly after GA induction, *P*_*θ*_ decayed, while *P*_*δ*_ increased, leading to a reliable band dominance change, a property that we use below for classifying distinct GA states. *P*_*δ*_ eventually decreased, while *P*_*θ*_ stayed low until the end of GA. We note that *P*_*δ*_ is weakly correlated to *p*_*t*_ (Fig. S1). Finally, the iso-electric suppression ratio tends to be higher during the incremented versus the other protocols (Fig. 3G). To conclude, the *θ*- and *δ*-bands are prominent during GA, with comparable behaviors across protocols.

### 3.3 Higher levels of IES precede the appearance of a *γ* pattern during recovery from GA

In some recordings, the EEG spectrogram during GA recovery revealed a stable and long-lasting activity in the *γ*-frequency domain (50−70 Hz range) (Fig. 4A). This phenomenon; which we refer to as *γ*-rebound, is characterized by a *γ*-power greater than that before and during GA (Methods section 5.10, Fig. 4A2,6).

**Figure 4:**
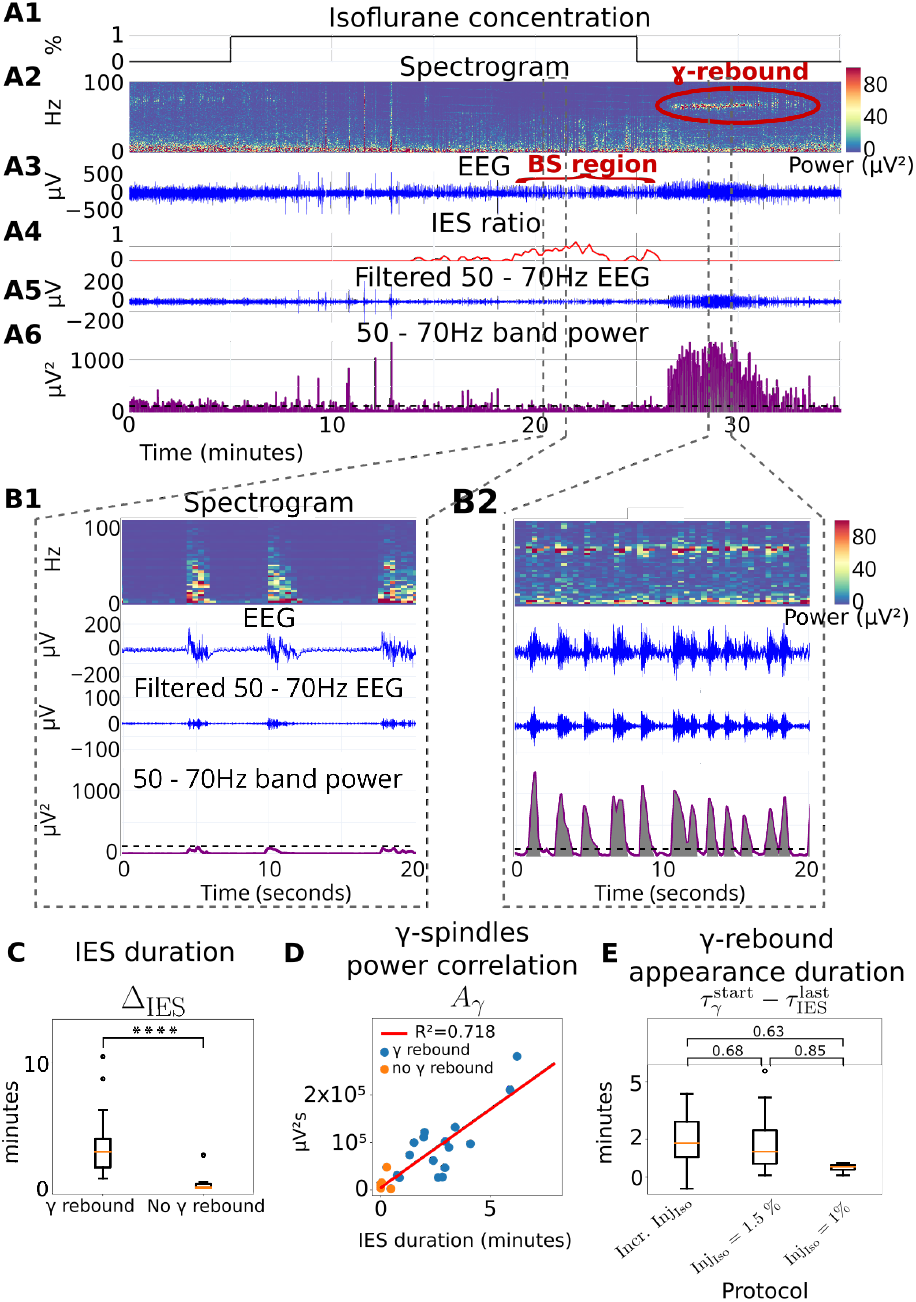
Cumulative IES duration *>* 30 seconds precedes *γ*-rebound during recovery from GA. **(A)** Spectral analysis of *γ*-rebound (red circle) that appear during recovery after burst suppression (BS region, red) characterized by IES ratio increase. *γ*-rebound shows significant amplitude in the 50 – 70 Hz power band, above threshold *T*_*γ*_ (black dashed lines). Magnification of 20-seconds plots **(B1)** during burst suppression and **(B2)** during *γ*rebound. **(C)** Statistics of IES duration in separated groups with and without gamma rebound, \*\*\*\**p <*0.0001 (two-sided Mann-Whitney U test). **(D)** Boxplot of *γ*-rebound appearance duration after the last IES given for the three anesthetic protocols and *p*-values of the associated two-sided Mann-Whitney U test. **(E)** Correlation between areas under *γ*-bursts power (grey area in B2) and IES durations.

To identify spectral features that would predict *γ*-rebound occurrence, we focused on IES. We noticed that *γ*-rebounds tended to appear after GA with burst suppression episodes (Fig. 4A3-4). Burst suppressions were identified visually as an alternation of IES and burst activity in the 0−50 Hz range [24] (Fig. 4B1). In parallel, the *γ*-rebound consisted of a succession of high amplitude and high power (purple) bursts located in a narrow frequency range around 60 Hz (Fig. 4B2).

To further characterize *γ*-rebound, we investigated whether it was associated with more time spent in IES. We found that *γ*-rebound was present in recordings where the cumulative IES duration (Δ_IES_) was greater than 30 seconds (Fig. 4C). Thus, recordings were differentiated into two groups (see Methods), those with and without *γ*-rebound, for which the cumulative IES duration was 198 *±* 144 seconds and 6.6 *±* 10.2 seconds respectively.

Finally, we investigated the relationship between the cumulative IES duration (Δ_IES_) and the power of the *γ*-rebound. To do so, we computed the instantaneous power in the 50*−*70 Hz range (*p*_*γ*_) (Fig. 4A6). We then computed the area (*A*_*γ*_) under the curve *p*_*γ*_ from the end of GA (time 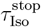) to the end of the recording (time 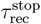). The distribution (Δ_IES_, *A*_*γ*_) (Fig. 4D) was fitted with a linear regression *y* = *ax* + *b*, where *a* = 32757*µV* ^2^, *b* = 6047*µV* ^2^*s*, and *R*^2^ = 0.72. This shows that the *γ*-burst power during recovery was highly correlated with the cumulative time spent in IES and thus mirrored depth of anesthesia. Additionally, we found no correlation between Δ_IES_ and the *γ*-rebound duration, or the *γ*-power *p*_*γ*_ (see Supplementary section S1.3 and Fig. S2).

We then studied the time of appearance of the *γ*-rebound relative to the last episode of IES. We found that the *γ*-rebound appeared a few minutes after the last IES (Fig. 4E), with no statistical difference across protocols. To conclude, we propose that *γ*-rebound is an *a posteriori* marker of long IES, which is characteristic of very deep anesthesia.

### 3.4 Transient sequence of EEG time-frequency patterns during isoflurane-induced GA

In this section, we present a segmentation of GA which incorporates the spectral decomposition done in section 3.2 and the *γ*-rebound described in section 3.3. To study whether EEG data during GA (Fig. 5A-B) could be segmented based on band tracking (Fig. 5C), power ratios (Fig. 5D), band suppressions (Fig. 5E), IES, and EEG and EMG spectrograms (Fig. 5F-G), we identified several key events which reliably happened in the same temporal order.

**Figure 5:**
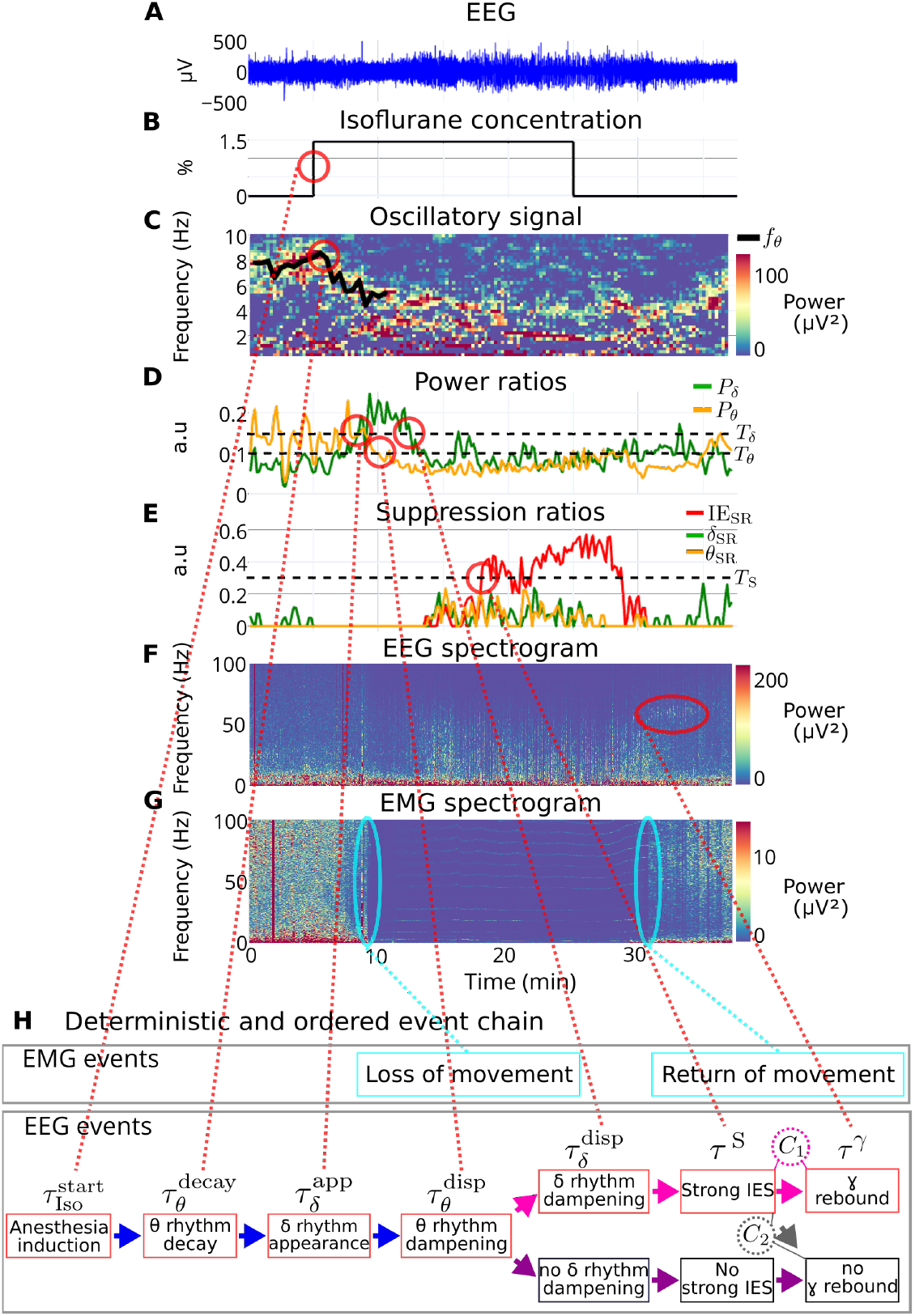
Ordered and deterministic chain of time-frequency events. **(A)** EEG recording. **(B)** Anesthesia protocol. **(C)** Extracted oscillatory signal and tracking of the center frequency *f*_*θ*_ of the *θ* rhythm. **(D)** Band power ratios. **(E)** Suppression ratios. **(F)** EEG spectrogram. **(G)** EMG spectrogram. **(H)** Deterministic and ordered frequency events chain. *C*_1_: cumulative IES time *>* 30 seconds. *C*_2_: cumulative IES time *<* 30 seconds.

The first relevant event was the beginning of GA, characterized by isoflurane delivery in the air (Fig H, first red box) at time 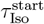. We then observed a decay of the *θ*-band, visible as a decrease of the *θ* center frequency *f*_*θ*_ (Fig. 5C). We refer to the beginning time of this phase as 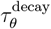 (eq. 16, Methods). Third, the *δ*-rhythm appeared, at tim 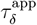 (eq. 9, Fig. 5D). Fourth and fifth, the *θ*-rhythm dampened at time 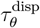 and the *δ*-rhythm dampened at time 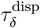 (eq. 10, Fig. 5D). Significant IES then emerged at time *τ* ^S^ (eq. 7, Fig. 5E). Finally, we observed the *γ*-rebound at time *τ*^*γ*^ (Methods section 5.10, Fig. 5G). The timestamps associated with these events are computed automatically (see Methods). In summary, the EEG events (Fig. 5H) appeared in the same temporal order across all recordings:

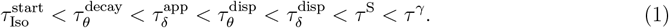

Finally, we quantified the duration between these EEG events and found that the order defined in equation 1 was valid on all recordings, although the duration between two consecutive events varied across protocols (Table 1). We conclude that in order to reach a specific event, the EEG first passes through all preceding states in a reproducible order. For instance, significant IES only occurred for recordings for which the *δ* band has dampened (Supplementary section 1.5 and Fig. S4).

**Table 1:** Main parameters.

| Parameter | Symbol | Value |
| --- | --- | --- |
| Sampling frequency (Hz) | $f_s$ | 500 Hz |
| Root Mean Square of the EEG | $\text{RMS}_{\text{EEG}}$ | |
| Relative IES threshold | $r_{\text{IES}}$ | 0.7 |
| Threshold for IES detection | $T_{\text{IES}} = r_{\text{IES}}\text{RMS}_{\text{EEG}}$ | |
| Threshold for $\theta$ suppressions detection | $T_{\theta\text{S}} = 0.2\text{RMS}_{\text{EEG}}$ | |
| Threshold for $\delta$ suppressions detection | $T_{\delta\text{S}} = 0.2\text{RMS}_{\text{EEG}}$ | |
| Suppression ratios time window width | 20 s |  |
| Suppression ratios time window overlap | 10 s |  |
| IES threshold | $T_S$ | 0.25 |
| Power ratio time window width |  | 20 s |
| Power ratio time window overlap |  | 0 s |
| Threshold for $\delta$ relative power | $T_\delta$ | 0.15 |
| Threshold for $\theta$ relative power | $T_\theta$ | 0.1 |
| Power ratio of the δ-band | $P_\delta(t)$ | |
| Power ratio of the θ-band | $P_\theta(t)$ | |
| Spectral time window width | $W_t$ | 60 s |
| Spectral time window overlap |  | 30 s |
| Coefficients of the 1/f-fit | $a_t, c_t$ | |
| Exponent of the 1/f-fit | $p_t$ | |
| Rhythm amplitude of the $\delta$ band | $b_\delta(t)$ | |
| Center frequency | $f_\delta(t)$ | |
| standard deviation | $\sigma_{\delta(t)}$ | |
| Rhythm amplitude of the $\theta$ band | $b_\theta(t)$ | |
| Center frequency | $f_\theta(t)$ | |
| Standard deviation | $\sigma_{\theta(t)}$ | |

**Table 2:**
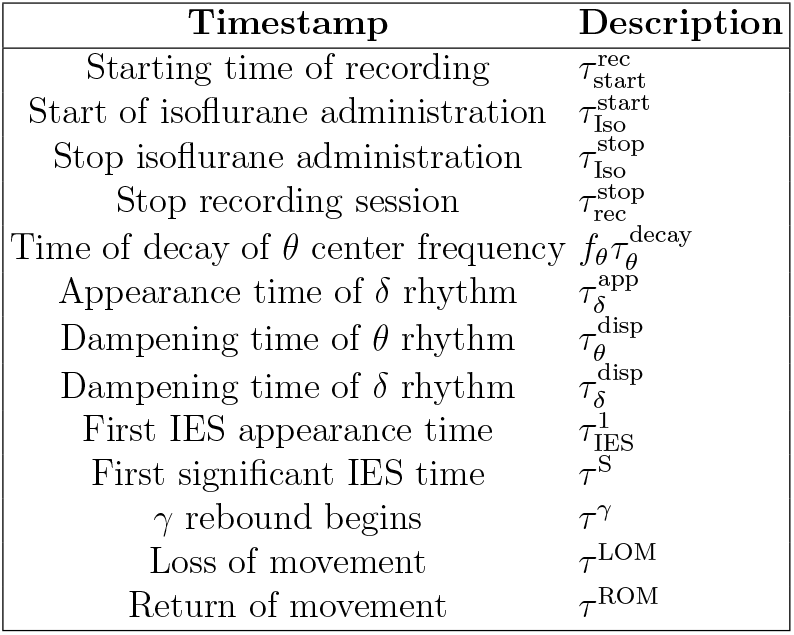
Timestamps of EEG and EMG time-frequency events.

We investigated the link between these EEG events and movement. Using the EMG, we identified the time of loss of movement 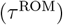 and the time of return of movement (*τ* ^ROM^) (Methods section 5.8, Fig. S10, and Fig. 5G). We found that the *θ*-rhythm dampening happened around the same time as LOM (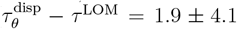 minutes). Likewise, when there was a *γ*-rebound, it started at the same time as ROM (*τ*^*γ*^ − *τ* ^ROM^ = 0.3 ± 0.8 minutes). However, the *γ*-rebound was not EMG contamination (Supplementary section S1.4 and Fig. S3).

To conclude, we report here a sequence of key, strictly ordered, events occurring during GA which are protocol-independent, with some variability in the transition durations between two consecutive events. This chain of events was characterized by the dynamics of the *δ*- and *θ*-bands. This sequence revealed two possible EEG behaviors: one path leading to long IES and *γ*-rebound, and another characterized by little or no IES and the absence of *γ*-rebound.

### 3.5 Predictive analysis and state-chart decomposition of isoflurane-induced GA

To further quantify the predictive value of the transient timestamps identified in section 3.4, we used a logistic regression (Methods section 5.9) to determine whether the delay to first IES occurrence 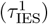, the delay to *θ*-band decay 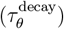, the delay to appearance of the *δ*-band 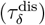, the delay to dampening of the *θ*-band 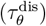, and the delay to dampening the *δ*-band 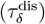 enable prediction of IES sensitivity. We define IES sensitivity as whether Δ_IES_ exceeds 30 seconds during GA, based on the *γ*-rebound analysis developed in section 3.3.

The univariate classification (Fig. 6A) logically revealed that the variable 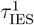 was most predictive of IES sensitivity, with a receiver operating characteristic area under the curve (ROC-AUC) value of 0.95. Interestingly, 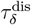 (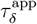 respectively) also carried a predictive power, characterized by a ROC-AUC value of 0.8 (0.65 respectively). The duration 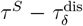 was 1.9 *±* 1.8 minutes for the incremented isoflurane protocol, and 4.5 ± 3.1 minutes for the 1.5% protocol (Supplementary section 1.5). Furthermore, the regression analysis showed that 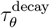 and 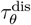 carried much less predictive power with ROC-AUC values of 0.4 and 0.36 respectively. We thus conclude that the *δ*-band dynamics carry more predictive value than the *θ*-band with respect to IES sensitivity.

**Figure 6:**
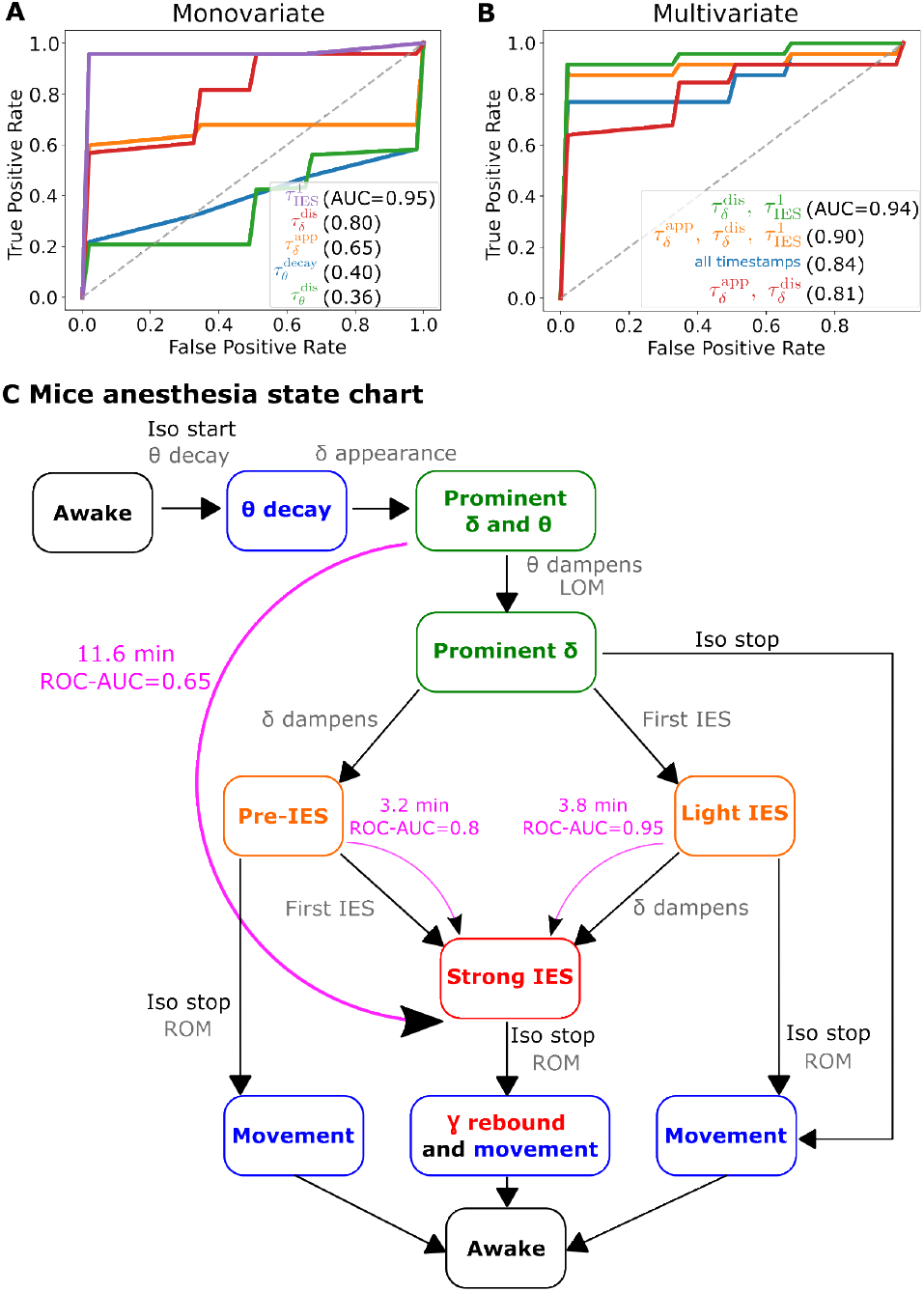
Logistic regression and state chart representation. IES Sensitive vs non-IES sensitive classification based on *θ, δ* and IES parameters. ROC curves and AUC are computed using a logistic regression classifier for **(A)** single predictors and **(B)** various combinations of predictors. **(C)** State chart characterizing the transitions between the different GA states starting from the awake state (black). Further states are associated with motion (blue), moderate depth of anesthesia (green), intermediate depth of anesthesia (orange) and high depth of anesthesia (red). They are characterized by the presence/absence of the *δ* and *θ* rhythms and IES. There are three main predictive states of IES sensitivity: the state “Prominent *δ* and *θ*” with a predictive power ROC-AUC = 0.65 and an average time delay to strong IES of 11.6 minutes, the state “Pre-IES” (ROC-AUC = 0.8 and average time of 3.2 minutes), and finally the state “Light IES” with a predictive power ROC-AUC = 0.95, and an average time of 3.8 minutes.

We then applied a multivariate logistic regression (Fig. 6B) to evaluate the predictive power of specific timestamps combinations. We found that the three models that performed 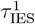: they were trained on the couple 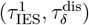 (ROC-AUC=0.94), on the triplet 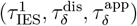 (ROC-AUC=0.9), or on all the timestamps (ROC-AUC=0.84). Finally, with the model trained only on *δ*-band appearance and dampening,we observed lower performance with ROC-AUC=0.81. From these analyses, we conclude that the most predictive variables are the first IES time 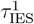, the *δ*-appearance time 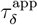, and the *δ*-disappearance time 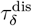. We also note that none of the multivariate models outperformed the univariate model trained on the first IES time 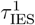 (ROC-AUC = 0.95).

To account for these deterministic relationships (Fig 5H and Table 1), we propose a state chart diagram as a synthetic graphical representation of the different states we previously reported: each state is characterized by *θ*- and *δ*-band spectral properties and suppression ratios (Fig. 6C). The initial state of the state chart, called “Awake”, is defined by a prominent *θ*-band with a stable *θ* center frequency *f*_*θ*_ (Methods eq 13), no prominent *δ*-band, and no IES. Subsequently, when isoflurane inhalation starts, the EEG switches to the second state called “*θ* decay”, in which *f*_*θ*_ decreases. The third state is “Prominent *δ* and *θ*”, where the *δ*-rhythm has appeared (Methods eq. 16). The fourth state is “Prominent *δ*”, where *θ* has dampened. Three states are accessible from there, which we detail below. In “Pre-IES”, the *θ*- and *δ*-bands are dampened and there is no IES. In “Light IES”, the *θ*-band is inactive, the *δ*-band is active, and there is little IES (0 IES_SR_ 0.25). The third state accessible from “Prominent *δ*” is called “Movement”, following ROM during recovery after isoflurane cessation. The next state after “Pre-IES” and “Light IES” is “Strong IES”, characterized by high IES_SR_ values (Methods eq. 7), and should be avoided. When GA stops, ROM happens, leading to the “*γ*-rebound and movement” state. If isoflurane stops in a different state than “Strong IES”, the EEG transitions to the “Movement” state without *γ*-rebound.

Interestingly, our statistics revealed an 11.6 minute delay between the states “Predominant *δ*” and “Strong IES”. This transition has a predictive value (ROC-AUC = 0.65). Moreover, the average transition time from the *δ*-rhythm disappearance state to the strong IES state is reduced to 3.2 minutes with a higher predictive (ROC-AUC = 0.8). Finally, the transition from the first IES time to the strong IES state occurs in 3.8 minutes on average, with a very high predictive value (ROC-AUC = 0.95).

In summary, we constructed a state chart associated with the depth of anesthesia, where each state is characterized by parameters computed from the EEG and EMG signals. Interestingly, the regression analysis revealed that the first IES time 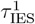, the *δ*-appearance time 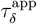, and the *δ*-dampening time 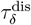 could be used to predict IES sensitivity and thus provide three different checkpoints that could be used in real-time analysis to monitor the brain IES sensitivity. These states were interlaced with clinical events observed during GA, namely GA induction, GA cessation, and loss and return of movement.

## 4 Discussion

Improved monitoring of depth of anesthesia is a crucial step to reduce post-operative complications. Beyond correcting anesthetic overdose, prevention of overdose is necessary to improve post-anesthetic outcome, but this has not been possible with current clinical EEG-based monitors. Our work here provides critical information that may make it more possible to predict and prevent overdose. We made a step forward in that direction. We developed a computational method based on EEG signal, and an automated and interpretable pipeline. Our approach revealed reproducible, timely ordered, brain states and transitions, that characterized the whole process of general anesthesia from induction to emergence. We further synthetize these states and transitions in a state-chart which represents depth of anesthesia in real-time and predicts IES onset several minutes in advance. This could prevent overdose, a task that has not been attained by the current clinical EEG-based monitors.

We conducted a time-frequency spectral analysis of EEG recorded during isoflurane-induced GA in mice. We decomposed the power spectrum density to extract prominent band activities, which turned out to be in the *θ*- and *δ*-domains during GA, and in the *θ*-, *δ*-, and *γ*-domains during recovery. We also computed the relative power of the *θ* and *δ* bands. We separated the oscillatory components from the 1/f-decay using the IRASA algorithm [23]. We tracked several frequency rhythms simultaneously and extracted the associated Gaussian parameters (Fig. 2). Finally, we detected IES using a threshold heuristic and computed the cumulative time spent in IES.

This method allowed us to identify a reliable chain of events (Fig. 5) which characterized the whole process of isoflurane anesthesia, fron induction to emergence. This led us to propose an EEG state chart that mirrors the whole duration and levels of anesthesia (Fig. 6C), although the underlying physiological mechanism remain unknown. This chart can be used to precisely titrate anesthetics and thus avoid undesirable states associated with high amounts of IES. Interestingly, we found that the *δ*-band dynamics were predictive of long IES several minutes in advance. Furthermore, although the *θ*-band was active during isoflurane GA, we found it had no predictive value regarding long IES. We hypothesized that the *θ*-band was linked to movement: the *θ*-band dampened around the same time as LOM. We also found that recovery from GA is not symmetric to induction: the EEG trajectory during recovery did not mirror the events chain of beginning and maintenance of GA. However, we observed during recovery a specific activity in the *γ*-band, namely the *γ*-rebound. Interestingly, this pattern appeared in recordings when the cumulative IES time was greater than 30 seconds. We therefore hypothesize that this pattern can be used as an *a posteriori* marker of very deep anesthesia.

### 4.1 Beyond the power spectral EEG decomposition

The EEG signal often mixes band oscillations with dominant 1/f-decay. Oscillatory bands present in the EEG can come from different brain regions, while the 1/f-decay is usually recognized as the background of spontaneous neuronal firing [25, 26, 11]. Decomposing the power spectral density of neurophysiological signals provides relevant insights on these signals [27]. It can be done by estimating the 1/fand oscillatory components directly on the power spectral density and fitting Gaussians to the oscillatory component, using the FOOOF algorithm [28]. Once the maximum amplitude frequency and band power are extracted, they can be used to quantify how *α*-oscillations and the 1/f-exponent are higher for young versus elderly patients [28]. However, this approach has several limitations. For instance, when applied to study the loss of consciousness (LOC) in humans anesthetized with propofol, it failed to detect oscillatory activity in the *δ*-domain [29]. This misclassification can be corrected using a convolution procedure on the EEG neuronal network [29].

Here, to overcome the low performance of the FOOOF algorithm in the low-frequency domain, we estimated the 1/f component using a different approach, namely IRASA [23], which uses successive re-samplings of the EEG signal directly, as opposed to the frequency domain. We then parameterized the remaining oscillatory component with Gaussians [30]. Interestingly, unlike previous approaches, our estimation dynamically tracks the time evolution of the *δ*- and *θ*-bands simultaneously (Fig. 2G). The algorithm we developed here allows for a robust and dynamic spectral decomposition that reveals the band characteristics during GA. This algorithm is fully automated and can be easily applied on other datasets.

### 4.2 Automated threshold selection for IES detection

In coma and GA, long IES episodes are associated with subsequent complications, such as confusion, delirium or loss of memory [31, 15]. These examples call for robust and automated IES detection in real-time. In most cases, long IES identification involves computing the fraction of time spent in IES in a sliding time window, (called the burst suppression ratio [32], similar to the present IES ratio IES_SR_), while the other fraction can contain any type of activity. Similarly, the burst suppression probability (BSP) [33] offers a binary segmentation for the presence of IES. Most algorithms [32, 33, 12] rely on a single-value threshold, where the value is fixed and not relative to the signal amplitude. This fixed threshold is bound to miss the interindividual variability of EEG amplitude [24]. A recent attempt to resolve this issue consists in estimating the variance of the EEG signal iteratively on successive time points [34], so that the threshold can be corrected by the variance of the signal up to time *t*. One limitation is that the model parameters need to be adjusted by an expert for each patient. Another direct approach for segmenting IES [13] is to compute the minimum between a fixed threshold value and the median of the difference between the upper and lower envelope of the signal, which accounts for individual signal variability. It however still includes a fixed threshold value.

Our data-driven approach is a step forward toward automatic and accurate IES detection. We propose a novel formula for IES threshold *T*_IES_ = *r*_IES_RMS_EEG_. The second coefficient accounts for inter-individual variability. The relative threshold *r*_IES_ was selected by continuously exploring detected IES duration with respect to *r*_IES_ (Supplementary section 2.1 and Fig. S6). This approach is robust on the present dataset, and it would be interesting to evaluate the goodness of fit of our optimal relative threshold value *r*_IES_ = 0.7 on a bigger dataset. The present heuristic could be adapted for real-time applications by evaluating the signal RMS(*t*) before the beginning of GA as opposed to during GA.

### 4.3 Revisiting the landmarks to control GA

The present approach extends the goal of monitoring depth of anesthesia beyond IES. Predicting IES occurrence in advance would help decrease the incidence of associated post-general anesthesia complications. This task is still difficult as we are missing a physiological model that could anticipate them based on the EEG signal. Recent works [12, 13] identified three parameters predictive of IES sensitivity in frontal EEG recordings of a human brain under propofol, namely the first appearance time of an *α* suppression, the slope of the *α*-suppression ratio, and the delay to the first IES occurrence. In contrast, in parietal EEG recordings of isoflurane-induced GA in mice, we observed that band suppressions were not predictive of IES sensitivity, because they did not reliably happen before IES. However, we did identify three parameters predictive of IES sensitivity: the appearance time of the *δ*-band, the dampening time of the *δ*-band, and the first IES time. The *δ* relative power seemed to be predictive here, analogous to the *α*-suppressions in humans with propofol. This difference may be due to the different drugs used [18], the different species recorded (human vs. rodent), or the different electrode locations.

### 4.4 Roles of *δ*- and *θ*-oscillations in isoflurane-induced GA

During isoflurane-induced GA of mice, we report oscillatory activity in the *δ*- and *θ*-frequency domains. These bands seemed to have distinct functions: while the *δ* dynamics accounted for neuronal responses to isoflurane and could anticipate IES, the *θ* dynamics seemed linked to mouse movement and had no statistical correlation with IES. Interestingly, following GA onset, the *δ*-rhythm appeared suddenly whereas the *θ*-rhythm decayed slowly.

It was shown that loss of consciousness is synchronized with an immediate *δ*-rhythm appearance in humans anesthetized with propofol [29], and high doses of sevoflurane induce coherent *δ*-oscillations in rats [14]. Our results also suggest that *δ*-oscillations reflect depth of anesthesia. In addition, we proved that, under fixed protocols, the time of appearance and disappearance of these oscillations are predictive of long IES.

*θ*-oscillations in rodent hippocampus have been linked to exploratory locomotion [35, 36]. However, the *θ*-oscillations we reported were observed in the cortex and are probably not from the hippocampus. Recent work has identified a neural rhythm called the respiration-entrained rhythm [37]. This rhythm is observable across several brain regions and peaks at the same frequency as breathing, around 12 Hz during exploration and 3 Hz during REM sleep. We hypothesize that part of the oscillatory activity reported here is this respiration-entrained rhythm. To conclude, the state chart (Fig. 6C) summarizes the complexity of the brain’s possible pathways during GA with isoflurane.

### 4.5 Characteristic *γ* activity during recovery from GA

A rebound activity in the *γ*-band appeared and persisted for several minutes after GA cessation (Fig. 4). This specific pattern only happened in recordings with cumulative IES duration greater than 30 seconds. Increased *γ* activity has been documented during awakening from 2 % isoflurane GA in rats, but no *γ*-rebound pattern was observed [38]. The isoflurane protocol used during recovery in that study [38] is very different from ours, which could explain why the *γ*-rebound was not observed then. We showed here that this *γ*-rebound was fully linked with the presence of long IES (Fig. 4E). We therefore hypothesize that the *γ*-rebound could reflect a network rebound, after a long period of hyperpolarization. The mechanisms underlying this manifestation remain to be clarified.

### 4.6 Potential clinical implications

Depth of anesthesia is routinely monitored in real-time via EEG during GA in human patients. These monitors process the EEG, and display an index between 0 and 100. Low index values indicate very deep sedation, while high index values indicate light sedation or wakefulness. Robust statistical associations were established between low EEG index values and poor outcome in large observational studies [39]. However, simple alerts on undesired EEG states have not been associated with improved outcome [40]. This could be because of the pharmaco-kinetic profiles of currently used hypnotic drugs. Indeed, once overdose is detected by the EEG monitor, its correction will require several minutes. This would imply that prevention and not only correction of the undesired EEG states would be necessary. Therefore, new methods to analyse and interpret the EEG states in routine clinical practice seem necessary.

We identified robust transition states between desired and undesired EEG states, and developed analytical tools to automatically detect these transition states in mice. We were indeed able to identify the transition states that precede IES, a typical undesired EEG state associated with poor outcome [41, 24]. Interestingly, this prediction is not possible with current EEG monitors for which IES episodes may be present even with index values within the desired range [42, 43]. Taken together, real-time identification of transitions could prevent these undesired states. Further studies are required in order to validate the robustness of these transition states in mice and humans who undergo GA. Finally, this approach needs to be further validated using other GA protocols in which the EEG and EMG states are paralleled with behavioral cues.

## 5 Methods

### 5.1 Animal care and use

All protocols were approved by the Institutional Animal Care and Use Committee at the University of California, San Francisco, and Gladstone Institutes, with institutional oversight. Experiments were conducted according to ARRIVE guidelines [44] and recommendations to facilitate transparent reporting [45]. All biological variables were documented. Adult C57BL/6J mice were used for each experiment. Mice of both sexes were used for the current study. Precautions were taken to minimize distress and the number of animals used in each set of experiments. Mice were housed in a pathogen-free barrier facility on a standard 12-hour light/dark cycle with ad libitum access to food and water.

### 5.2 Surgical implantation of the EEG and EMG devices and EEG recordings

Seventeen adult mice underwent surgical implantation of EMG and EEG devices for chronic electromyogram and electrocorticogram recordings. Mice were anesthetized with vaporized isoflurane (3% induction, 1 – 2% maintenance, carried by 100% *O*_2_ at a flow rate of 2 L/minute) and placed under a stereotaxic frame for chronic EEG implants as previously described in [46, 47]. Briefly, an EEG screw was implanted in the skull overlying the cortical region at the following coordinates: 1.0 mm anterior from Bregma and 2.5 mm lateral from the midline [48]. A ground screw was placed overlying the cerebellum (0.5–1 mm posterior to Lambda and 0.5–1 mm lateral to midline). The EMG electrode was placed in the deep parasagittal cervical muscles. The skin was closed over the entire apparatus, which was sealed with dental cement and Vetbond tissue adhesive. For analgesia, topical lidocaine ointment (5%) was applied prior to incision and extended-release buprenorphine (0.05–0.1 mg/kg s.c.) was administered prior to recovery from anesthesia. Mice recovered for 5–7 days after surgery before the start of recordings.

Two types of EEG devices were used: purchased wireless telemetry devices (HD-X02, Data Sciences International (DSI), St. Paul, MN) and custom-made wired EEG devices made in the Paz lab using cortical screws connected to a Millmax device [47]. Wireless recordings were acquired using Ponemah software (DSI); wired recordings were acquired using Synapse software (Tucker Davis Technologies).

### 5.3 Protocols using isoflurane-induced GA

GA was induced in the mobile isoflurane anesthesia induction chambers. Mice were placed into the plastic chambers, and EEG was recorded for 5 minutes before vaporized isoflurane carried by 100% O2 at a flow rate of 2L/min was turned on. There were three different protocols for isoflurane-induced GA. In the first, the isoflurane dose was gradually increased from 0.5% to 2% every 5 minutes with 0.5% increments (n = 14), while in the other two, the dose was fixed at 1% (n = 10) and 1.5% (n = 9) (Fig. 1A-B). Vaporized isoflurane was turned off after 20 minutes. Mice were kept in the plastic induction chamber until full recovery and visible unimpaired movement around the plastic chamber, usually no more than 20 minutes. Mice were returned to home cages at the end of the experiment. These protocols were chosen in order to characterize and compare the mouse EEG response to isoflurane under constant light sedation (1% protocol), constant high sedation (1.5% protocol), and increasing concentration from light to high sedation (incremented protocol). Some mice were used for several protocols, with at least one resting day between two sessions. Eight mice underwent the incremented and 1.5% protocol, four mice underwent the step protocol alone, three mice underwent the 1% protocol, and two mice underwent the 1% protocol three time, the 1.5% protocol once, and the incremented protocol once.

### 5.4 EEG data and pre-processing

The EEG signal *S*(*t*) was digitized at a sampling frequency *f*_*s*_ = 500 Hz. We first identified artifacts, like regions where the signal is constant and equal to 0 due to no signal being recorded. Regions with abnormally high values were also identified as artifacts using hysteresis thresholding with a low threshold of 0.08 *µV* and a high threshold of 1200 *µV* [49]. We labeled the artifact regions as NaN (not containing any significant signal to be processed). Three EEG recordings (one per anesthesia protocol) were excluded because they contained too many artifacts.

### 5.5 Signal processing tools

Signal processing notations used throughout the Methods section are defined here. For some computations specified below,the signal *S* was band-pass filtered using a Butterworth forwardbackward filter [50] of order 1 (effective order 4) in a frequency domain [*f*_1_, *f*_2_], where the frequencies *f*_1_ and *f*_2_ are specified in Table 1. A sliding window *W*_*w*_(*t*) centered at time *t* and of width *w* was used to compute statistical markers locally in time. For *w ∈* ℝ^+^ and 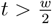, we use the notation

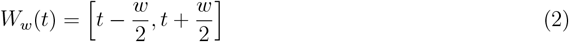

We used an overlap between two sliding windows of 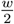 or 0. Chosen values of *w* and overlap are specified in Table 1.

#### 5.5.1 EEG segmentation of suppression periods

To detect IES, regions where the amplitude of the EEG was smaller than the threshold *T*_IES_ for at least 1 second were identified. The threshold chosen was

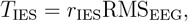

where *r*_IES_ = 0.7 and RMS_EEG_ is the Root Mean Square of the entire recording:

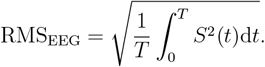

The threshold was chosen using a novel heuristic, see supplementary section 2.1, Fig. S5, and Fig. S6. Two detected suppressions separated by less than 0.5 s were merged as one single suppression [12]. The detection of IES is shown in Fig. 1C1.

Similarly, suppressions of the *θ*-(resp. *δ*-) rhythm were tracked. The signal was band-pass filtered within the 5*−*10 Hz (resp. 2.5*−*4.5 Hz) band. The segments where the amplitude of the filtered signal is below the threshold *T*_*θ*S_ = 0.2 RMS_EEG_ (resp. *T*_*δ*S_ = 0.2 RMS_EEG_) were labeled as *θ*-(resp. *δ*-) suppressions (Fig. 1C2-3). IES can also be detected as *θ* suppressions and *δ* suppressions. Therefore, segments detected as IES are removed from *θ* and *δ* suppressions.

#### 5.5.2 Estimating suppression ratios

We define the iso-electric suppression Ratio (IES_SR_) as the proportion of time that the EEG signal spends in IES inside a sliding window *W*_*R*_(*t*) of width *R* (eq. 2), that is

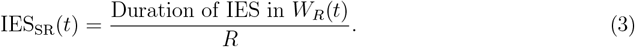

Similarly, we define the *θ*-Suppression Ratio *θ*_SR_ as

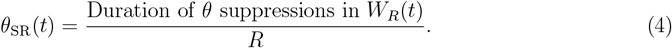

Finally, the *δ*-Suppression Ratio is

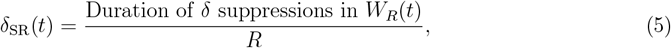

These suppression ratios are computed on sliding time windows of width *R* =20 seconds and an overlap of 10 seconds (Fig. 1E). Notably, the iso-electric suppression ratio is very similar to the commonly used burst suppression ratio [32].

#### 5.5.3 Detecting strong IES episodes

Segments where the IES ratio (eq. 3) was high were collected. IES at time *t* is considered strong if more than a threshold *T*_*S*_ = 25% of the time window centered in *t* is detected as IES:

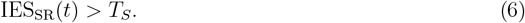

This threshold was chosen empirically after a visual inspection of EEG.

Therefore, the first strong IES time *τ* ^S^ is the first time at which IES_SR_(*t*) *> T*_*S*_ on consecutive time windows for at least 40 seconds.

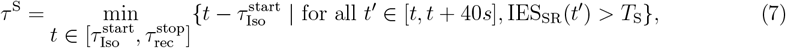

where 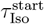 is the beginning of induction time, and 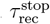 is the end of recording time.

### 5.6 Computing frequency power ratios from EEG and detecting present frequency rhythms

#### 5.6.1 Computing power ratios of the *δ*- and *θ*-bands

The power of the low-pass filtered EEG *S*_20_ under 20 Hz, the band-pass filtered signal *S*_*θ*_ in 5 *−* 10 Hz, and *S*_*δ*_ band-pass filtered in 2.5 *−* 4.5 Hz were computed. The associated powers are computed in the sliding time window *W*_*R*_(*t*):

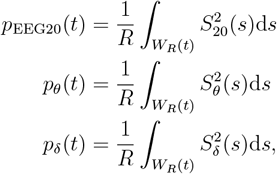

The *θ*- and *δ*-power ratios are defined by

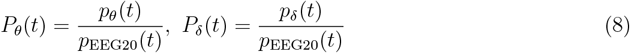

and computed over a sliding windows *W*_*R*_(*t*) with *R* = 20 seconds and no overlap (eq. 2, fig. 1E).

#### 5.6.2 Activity of frequency bands

To assess whether the *θ*- or *δ*-rhythm is prominent, its power ratio is computed (eq. 8) over the sliding windows defined above (eq. 2) and compare it to a threshold *T*_*θ*_ = 0.1 or *T*_*δ*_ = 0.15. The *θ* rhythm (respectively *δ* rhythm) is considered to be prominent at time *t* if *P*_*θ*_ *> T*_*θ*_ (respectively *P*_*δ*_ *> T*_*δ*_) for at least 1 minute. Conversely,the dampening time of the *θ* rhythm (respectively *δ* rhythm) is defined as when *P*_*θ*_ *< T*_*θ*_ (respectively *P*_*δ*_ *< T*_*δ*_) for at least 1 minute. For instance in Fig. 1E, the *δ* rhythm is absent during the 0 *−* 11 minutes period, prominent during the 11 *−* 22 minutes period, dampened during the 22 − 27 minutes period, and prominent during the 27 *−* 33 minutes period. The first time of *δ* appearance 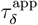 is defined as the first time at which the *δ* rhythm is prominent since the beginning of anesthesia:

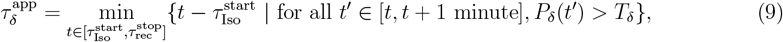

where 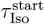 is the beginning of induction time and 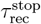 is the end time of the recording. The first time of *θ* appearance is not defined, because this rhythm is already prominent before the beginning of anesthesia. Similarly, the time of dampening of the *θ* and *δ* rhythm are defined as follows:

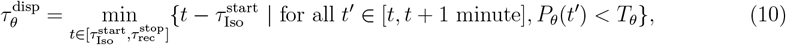

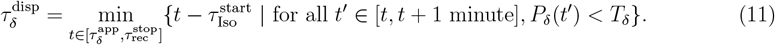

### 5.7 Extracting dominant frequency rhythms present in the EEG

To quantify the persistence of a frequency rhythm present in the EEG signal,an algorithm [23] which separates the power spectrum into 1*/f* and oscillatory components was adapted. The novelty consists in using this decomposition in sliding time windows, which results in a decomposition that is continuous over time.

#### 5.7.1 Spectral decomposition on a sliding time window

The power spectral density PSD_*t,w*_(*f*) of the signal *S* is computed over the time window *W*_*w*_(*t*) (eq.2) and can be decomposed as follows:

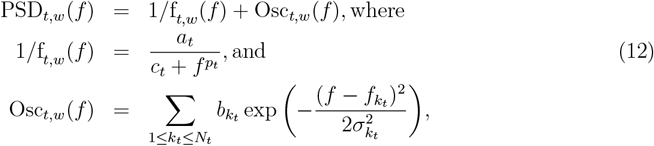

see Fig. 2A and [11]. The first component 1*/f*_*t,w*_ captures the frequency decay of the power spectral density and is characterized by the amplitude *a*_*t*_, the exponent *p*_*t*_ and the correction term *c*_*t*_, which we estimate, as discussed below. *c*_*t*_ was used for two reasons. First, the signal power spectral density does not diverge in the small frequency domain (Fig. 2A2). *c*_*t*_ *>* 0 corrects this modelization error. Second, introducing *c*_*t*_ divides by 15 the approximation error, see supplementary section S2.2, Fig. S8 and Fig. S9. The second component Osc_*t,w*_ accounts for the oscillatory part of the signal and can be decomposed as a sum of *N*_*t*_ Gaussians (Fig. 2A3-4). Each Gaussian peaks at the main frequency 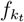 with a standard deviation 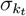 and amplitude 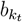. The oscillatory parameters to be estimated are the number *N*_*t*_ and the Gaussian parameters 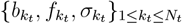. In practice, there are at most two main components in the 0*−*15 Hz domain: the *θ* and *δ* bands, so that

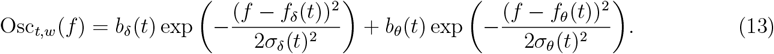

#### 5.7.2 Parameters estimation on a time window

Estimation of the 1/f and oscillatory parameters of the EEG signal *S* over the time window *W*_*w*_(*t*) was done by a fully automated algorithm using the steps described below.

1. The power spectral density of the signal *S* was computed in the range 0.2*−*15 Hz the power spectral density of the signal *S* (Fig. S7B) over *W*_*w*_(*t*) using Welch’s method [51] (with a 5 seconds sub-window).
2. The IRASA method [23] was used to estimate the 1/f component of the power spectral density. It consists of applying several scaling factors *h* on the signal *S*, computing the corresponding power spectral densities, and the median of the power spectral densities provides the 1*/f* component. In practice, the signal *S* in *W*_*w*_(*t*) was up-scaled and downscaled using factors *h*_*i*_ between 1.1 and 1.9 with a 0.05 increment and their reciprocals 1*/h*_*i*_. Then, the geometrical mean PSD_*GM*(*i*)_ of 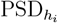 and 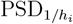 was computed for each i. Finally, the 1*/f* estimate was the median PSD_*m*_ of the PSD_*GM*(*i*)_ for all i (Fig. S7C, light blue curve). The parameters *a*_*t*_, *p*_*t*_, and *c*_*t*_ were obtained by fitting PSD_*m*_ on the 0.2*−*15 Hz interval in the log-log scale (Fig. S7C, red curve). The YASA python library [52]was used to implement the IRASA method, and the SciPy Python module [53] to fit parameters *a*_*t*_, *p*_*t*_, and *c*_*t*_.
3. The oscillatory component was computed by removing the estimated 1/f component from the power spectral density (Fig. S7D):

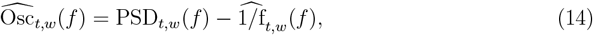

where 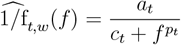.
4. The oscillatory parameters *N*_*t*_ and 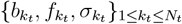 were estimated by fitting a sum of Gaussian functions to the oscillatory component 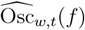 on the frequency domain 1*−*15 Hz. This part is a generalization of a method developed in [30].
5. Gaussian components where the standard deviation 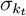 was either smaller than 0.2 or larger than 2 were discarded. In addition, Gaussian components with too small amplitude 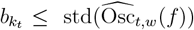 were discarded. The thresholds 0.2 and 2 were chosen empirically and the threshold std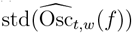 was inspired by [28].
6. At most one Gaussian was selected for which center frequency 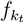 falls into the *θ* (resp. *δ*) 4*−*10 Hz (resp. 2*−*4 Hz) band (Fig. S7E). When there were several detected Gaussians that fell into one frequency band, the Gaussian component with the largest area 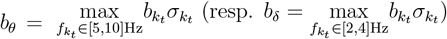 was selected.

When no Gaussian component was present in one of the band domains, we considered that there was no prominent rhythm in this band for this time window. In the rare cases where two Gaussians had the same maximum area, we selected the one for which the center frequency 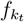 was closest to the median frequency, i.e 7.5 Hz for the *θ*-band, 3 Hz for the *δ*-band.

#### 5.7.3 Rhythm tracking along GA

Subsection 5.7.2 estimated the Gaussian *G*_*θ*_(*f, t*) (resp. *G*_*δ*_(*f, t*)) of the *θ*-(resp. *δ*-) rhythm in one time window of width *w* = 60s and centered at time *t*. To define a continuous estimation for the time-varying *θ* and *δ* rhythms, the windows *W*_*w*_(*t*_*i*_) at discretized times *t*_*i*_ with a 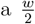 step are used. To define a continuous curve, from these discretized windows *W*_*w*_(*t*_*i*_), the band Gaussian on *W*_*w*_(*t*_*i*_), *i* ∈ ℕ is estimated as described in subsection 5.7.2.

When a Gaussian is detected in the consecutive frames *W*_*w*_(*t*_*i*_) and *W*_*w*_(*t*_*i*+1_), it is interpolated by a straight line the center frequencies (*f*_band_(*t*_*i*_), *f*_band_(*t*_*i*+1_)) in the time interval [*t*_*i*_, *t*_*i*+1_]. The same procedure is also applied to the standard deviation. However, when a band Gaussian is detected in *W*_*w*_(*t*_*i*_) but none are detected in either *W*_*w*_(*t*_*i−*1_) or *W*_*w*_(*t*_*i*+1_), it is not interpolated and the band rhythm is considered not significant.

These steps are applied for both the *θ* and *δ* bands. Finally, to further quantify the dynamics of each rhythm,the dynamics of the Gaussian width were followed. The lower and upper curves centered at the frequency *f* (*t*) of the fitted Gaussian, each at a standard deviation *σ*(*t*) distance were applied so that

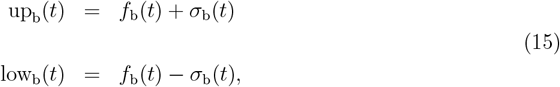

where *b* ∈ {*θ, δ*}. Thus the distance between the two curves is precisely twice the variance up_b_(*t*) low_b_(*t*) = 2*σ*_b_(*t*). See Fig. 2G for an example of continuous rhythm tracking, with *w* =1 minute.

#### 5.7.4 Decay of *θ* center frequency during induction

A common behavior of the *θ* rhythm was observed in all recordings. At baseline, the *θ* rhythm is prominent, and its center frequency *f*_*θ*_ is stable around 8 Hz. Then, shortly after the beginning of anesthesia, *f*_*θ*_ decreases rapidly for several minutes (Fig. 2). The start time of the *f*_*θ*_ decay 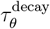 is defined as:

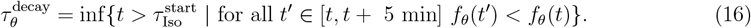

### 5.8 Identifying loss and return of movement from EMG

Loss and return of movement were identified by visually inspecting the EMG (Fig. S10). Loss of movement is easily recognizable on the EMG as a switch from active to flat signal. Similarly, the return of movement is a switch from flat to active signal. Notably, the EMG recordings contain artifacts, probably from respiration and electrocardiogram. These artifacts were of very low amplitude and therefore did not impact LOM and ROM identification.

### 5.9 Logistic regression analysis

A logistic regression approach with *l*_2_ regularization [54] and a regularization constant C=1 were used to evaluate the contribution of several parameters to the prediction of significant time spent in IES. The criterion for class separation was chosen with the *γ* rebound phenomena, which happens for recordings with more than 30 seconds in IES (section 3.3). The dataset was thus divided into a positive class (total time spent in IES is more than 30 seconds) and a negative class (total time spent in IES is less than 30 seconds).

The dataset was separated into training and validation sets using a stratified group k-fold strategy: this strategy ensures that recordings from one individual are not split between the train and validation set (group) and that the validation sets of all folds have equivalent proportions of positive and negative labels (stratified). The scikit-learn python library [55] was used with a 4 folds separation, due to the limited number of recordings in the negative class.

The logistic regression models were fed timestamps of events identified in section 3.4. Due to constraints on the various durations, each time feature was normalized given to the model in the following way:

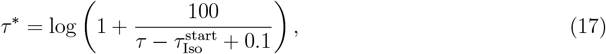

where *τ* is the time feature computed by our algorithm, *τ* is the feature fed to the logistic regression model, and 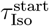 is the time of the beginning of anesthesia. This choice was inspired by [13].

### 5.10 *γ*-rebound identification

The *γ*-rebound recordings were identified when the *γ*-power was significantly larger during recovery than in the rest of the recording for at least 2 minutes, using a threshold *T*_*γ*_ on the power defined by

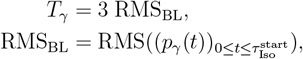

where *p*_*γ*_(*t*) is the power of the filtered 50−70 Hz EEG signal, computed on sliding windows of width 0.2 seconds and zero overlap and 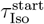 is the instant where the isoflurane starts to be administered.

## Acknowledgments

D.H. research is supported by a grant ANR NEUC-0001, CNRS pre-maturation, and the European Research Council (ERC) under the European Union’s Horizon 2020 research and innovation program (grant agreement No 882673). Experimental work in the J.T.P. lab was supported by Gladstone Institutes. We thank Kathryn Clairborn, Christophe Sun, and Matteo Dora for critical feedback on our manuscript.

## Authors contributions

Y.V., B.L., and D.N. all contributed equally to data collection, and editing of the manuscript. V.L. wrote the code. V.L. and D.H. analyzed the data. V.L. and D.H. wrote the manuscript. D.H. and J.T.P. supervised and designed the research. D.L. and J.T.P edited the manuscript.

